# *Atypical Chemokine Receptor 1 (Ackr1)*-deficient Mice Resist Lethal SARS-CoV-2 Challenge

**DOI:** 10.1101/2023.06.05.543759

**Authors:** Shamik Majumdar, Joseph D. Weaver, Sergio M. Pontejo, Mahnaz Minai, Shalamar Georgia-Clark, Ji-Liang Gao, Xinping Lu, Hongwei Zhang, Nicole Lackemeyer, Tracey Burdette, Reed Johnson, Brian L. Kelsall, Joshua M. Farber, Derron A. Alves, Philip M. Murphy

**Affiliations:** Molecular Signaling Section, Laboratory of Molecular Immunology, National Institute of Allergy and Infectious Diseases, National Institutes of Health, Bethesda, MD, USA; Infectious Disease Pathogenesis Section, Comparative Medicine Branch, National Institute of Allergy and Infectious Diseases, National Institutes of Health, Bethesda, MD, USA; SARS-CoV-2 Virology Core, Laboratory of Viral Diseases, National Institute of Allergy and Infectious Diseases, National Institutes of Health, Bethesda, MD, USA; Inflammation Biology Section, Laboratory of Molecular Immunology, National Institute of Allergy and Infectious Diseases, National Institutes of Health, Bethesda, MD, USA; Mucosal Immunobiology Section, National Institute of Allergy and Infectious Diseases, National Institutes of Health, Bethesda, MD, USA

**Keywords:** COVID-19, endothelium, inflammation, mouse

## Abstract

High pro-inflammatory chemokine levels have been reported in blood and lung in patients with COVID-19. To investigate specific roles in pathogenesis, we studied the regulation of chemokine ligands and receptors in the lungs of 4-6-month-old wild type C57BL/6 mice infected with the MA10 mouse-adapted strain of SARS-CoV-2. We found that atypical chemokine receptor 1 (*Ackr1*, also known as Duffy antigen receptor for chemokines/DARC) was the most highly upregulated chemokine receptor in infected lung, where it localized to endothelial cells of veins and arterioles. In a screen of 7 leukocyte chemoattractant or chemoattractant receptor knockout mouse lines, *Ackr1*^-/-^ mice were unique in having lower mortality after SARS-CoV-2 infection, particularly in males. ACKR1 is a non-signaling chemokine receptor that in addition to endothelium is also expressed on erythrocytes and Purkinje cells of the cerebellum. It binds promiscuously to both inflammatory CC and CXC chemokines and has been reported to control chemokine availability which may influence the shape of chemotactic gradients and the ability of leukocytes to extravasate and produce immunopathology. Of note, erythrocyte ACKR1 deficiency is fixed in sub-Saharan African populations where COVID-19 has been reported to result in low mortality compared to worldwide data. Our data suggest the possibility of a causal contribution of ACKR1 deficiency to low sub-Saharan COVID-19 mortality and identify ACKR1 as a possible drug target in the disease.

## Introduction

Since its emergence in humans (Zhu et al., 2020), severe acute respiratory syndrome coronavirus-2 (SARS-CoV-2), the causative agent of the coronavirus disease-2019 (COVID-19) pandemic, has resulted in at least 6.8 million deaths (https://coronavirus.jhu.edu/map.html). After more than three years and despite widespread prior infection and vaccination, SARS-CoV-2 still claims over 1000 deaths weekly in the United States (https://covid.cdc.gov/covid-data-tracker/#datatracker-home) and is the leading infectious and respiratory cause of death among children (Flaxman et al., 2023).

After inhaling SARS-CoV-2-contaminated respiratory droplets and aerosols, the clinical outcome is highly heterogeneous, ranging from asymptomatic infection to life-threatening illness, including acute respiratory distress syndrome (ARDS) (Wang et al., 2020; Acute Respiratory Distress Syndrome: The Berlin Definition, 2012). In a retrospective study of 201 patients with COVID-19 pneumonia, 52.4% of patients who developed ARDS died (Wu et al., 2020). While lung pathology included microvascular clotting and prominent myeloid cell infiltration (Zaid et al., 2021; Lamers and Haagmans, 2022), the molecular mechanisms have not been fully defined.

Broncho-alveolar lavage fluid (BALF) from patients with severe COVID-19 contain monocytes and macrophages that have been reported to produce extremely high amounts of cytokines and chemokines, which likely initiate and potentiate cytokine storm, leading to ARDS (Merad and Martin, 2020; Xu et al., 2020). Levels of serum and BALF chemokines differ depending on the severity of COVID-19 in humans (Majumdar and Murphy, 2021). To assess the functional importance of chemokines and their receptors experimentally, we analyzed a mouse model of SARS-CoV-2 infection, reasoning that any receptor that promoted severe disease outcome might provide a host target for anti-inflammatory drug development.

## Materials and methods

### Mice

JAX664 WT C57BL/6J mice were purchased from The Jackson Laboratory (Bar Harbor, ME, USA). The following genetically modified mice were used in the study: *Acrk1*^-/-^ (*Ackr1*^tm1Scp^/*Mmnc*, gift of Stephen Peiper) (Dawson et al., 2000), *Ackr2*^-/-^ (B6.129/Sv-Ccbp2^tm1Gjg^/Apb, gift of Gerard Graham) (Jamieson et al., 2005), *Camp*^-/-^ (Chen et al., 2014) and *Fpr1/2*^-/-^ (Liu et al., 2012) (gift of Ji Ming Wang), *Ccr5*^-/-^ (B6.129P2-Ccr5^tm1Kuz^/J, The Jackson Laboratory) (Kuziel et al., 2003) and *Cxcr3*^-/-^ (B6.129P2-*Cxcr3tm1*/J, The Jackson Laboratory). *Ccr6*^-/-^ mice were generated by Ozgene (Perth, WA, Australia) by crossing B6.Ccr6^fl/fl^Oz mice with CMV-cre mice.

The mice were maintained and bred under specific pathogen-free conditions with access to food and water *ad libitum*. Mice at 4-6 months of age were used for infection experiments. The studies were performed in accordance with NIH institutional guidelines and were conducted in Assessment and Accreditation for Laboratory Animal Care-accredited Biosafety Level (BSL) 2 and 3 facilities at the NIAID/NIH using a protocol approved by the NIAID Animal Care and Use Committee.

### Virus stock preparation and mouse infections

Mouse-adapted (MA) SARS-CoV-2 was obtained from the Biodefense and Emerging Infections Research Resources Repository (BEI resources, Manassas, VA, USA). Stocks were generated by infecting Vero E6 cells expressing TMRPSS2 (Liu et al., 2021) at a multiplicity of infection of 0.01, incubated for 48 h at 37°C in 5% CO_2_. Media containing virus was harvested, centrifuged to remove cellular debris, and aliquoted. Stocks were sequenced to confirm single nucleotide variants (SNVs) consistent with MA SARS-CoV-2. Stock generation was performed in a BSL-3 containment laboratory.

All SARS-CoV-2 infections were performed within an animal BSL-3 containment laboratory. Mice were anesthetized by isoflurane inhalation, and a dose of 8.62 × 10^4^ TCID_50_ SVG-025 was administered by intranasal instillation to anesthetized mice. Body weights of mice were recorded on the day of infection, and until day 14. The mice were monitored for morbidity and were euthanized per protocol if the body weight dropped by ≥ 25% of the initial body weight.

For H1N1 Influenza A virus (IAV, A/Puerto Rico/8/1934, PR8) infections, the virus stocks were provided by Dr. Jonathan W. Yewdell (National Institute of Allergy and Infectious Diseases). One hundred TCID_50_ of IAV were administered by intranasal instillation to isoflurane-anesthetized mice under BSL-2 containment. The IAV- and mock-infected mice were euthanized on day 4 post-infection and lungs were processed for RT-PCR assays.

Mouse cytomegalovirus (MCMV) stocks were generated from a bacmid for the reporter MCMV-3DR strain (kindly provided by Martin Messerle, Hannover University) as previously described (Marquardt et al., 2011; Stahl et al., 2015). Virus was purified by ultracentrifugation from the supernatant of infected M2-10B4 cells (ATCC, Gaithersburg, MD) maintained in RPMI-Glutamax media (Thermo Fisher Scientific, Waltham, MA) supplemented with 10% FBS at 37°C in 5% CO_2_. Viral titers were determined by plaque assay in M2-10B4 cells and stocks were stored at -80°C in virus standard buffer (VSB, 50 mM Tris-HCl pH 7.8, 12 mM KCl and 5 mM EDTA). About 10^5^ pfu of MCMV in PBS were administered by intranasal instillation to isoflurane-anesthetized mice under BSL-2 containment. The MCMV- and mock-infected mice were euthanized by cervical dislocation 4 days post-infection (dpi) and lungs were dissected and homogenized in PBS using an Omni tissue homogenizer (Omni International, Kennesaw, GA).

### Chemokine and cytokine analysis by qPCR

Lungs were homogenized in 1 ml of PBS or HBSS (Thermo Fisher Scientific) and 100 µl of the homogenate were added to 1:3 v/v of TRIzol™ LS (Thermo Fisher Scientific) and incubated for 10 min at room temperature to inactivate the virus. After extraction with chloroform, the aqueous phase was mixed 1:1 v/v with 70% ethanol and loaded into a column of the Isolate II RNA Mini kit (Bioline, Cincinnati, OH). Total RNA was purified following the manufacturer’s recommendations, which included a step to remove genomic DNA. cDNA was synthetized from 200 ng of RNA using the SensiFAST cDNA Synthesis kit (Bioline) and gene expression was analyzed by qPCR in a CFX96 Touch Real-Time PCR instrument (Bio-Rad, Hercules, CA) using the SensiFAST SYBR Hi-ROX kit (Bioline). Predesigned KiCqStart SYBR green primers for the analyzed cytokines, chemokines, chemokine receptors, and reference genes were acquired from MilliporeSigma (Burlington, MA). The expression fold change over the mock-infected group for each gene was calculated as 2EXP(-ΔΔCt) using the geometric mean Ct of 3 genes (*Gapdh*, *ActB* and *Hprt1*) as reference.

### Histopathology

Following tissue collection under BSL-3 conditions, harvested lungs from SARS-CoV-2 infected and mock-infected mice were immersed and fixed in 10% neutral buffered formalin (MilliporeSigma) for a minimum of 3 days, and then transferred to 70% ethanol. Lung samples were processed per a standard protocol. Briefly, tissues were processed with Leica ASP6025 tissue processor (Leica Microsystems, Wetzlar, Germany), embedded in paraffin, and sectioned at 5 μm for histological analysis. Tissue sections were stained with hematoxylin and eosin (H&E) for routine histopathology and then evaluated by a pathologist in a blinded manner. Sections were examined by a board-certified veterinary pathologist (DAA) using an Olympus BX51 light microscope and photomicrographs were taken using an Olympus DP28 camera.

### Immunohistochemistry

Formalin-fixed paraffin-embedded (FFPE) tissue sections (5 µm) obtained from SARS-CoV-2 infected and mock-infected mouse lung tissues were used to perform immunohistochemical (IHC) staining using the following antibodies: a rabbit polyclonal anti-CD31 (Abcam, Cambridge, United Kingdom, Catalog No: ab28364, dilution of 1:50) and a rabbit polyclonal anti-SARS-CoV-2 (GeneTex, Irvine, CA, Catalog No: GTX135357, dilution of 1:2000). Staining was carried out on the Bond RX (Leica Biosystems, Buffalo Grove, IL) platform according to manufacturer-supplied protocols. Briefly, 5 µm sections were deparaffinized and rehydrated. Heat-induced epitope retrieval (HIER) was performed using Epitope Retrieval Solution 1, pH 6.0, heated to 100°C for 20 min. The specimen was then incubated with hydrogen peroxide to quench endogenous peroxidase activity prior to applying the primary antibody. Detection with DAB chromogen was completed using the Bond Polymer Refine Detection kit (Leica Biosystems, Catalog # DS9800). Slides were finally cleared through graded alcohol and xylene washes prior to mounting and placing cover slips.

### RNAScope® *in situ* Hybridization (ISH)

RNA ISH was carried out on the Bond RX auto-Stainer (Leica Biosystems) platform according to manufacturer-supplied protocols using the Bond Polymer Refine Red Detection kit (Leica Biosystems) along with the RNAscope**®** 2.5 LS Reagent kit-Red (Advanced Cell Diagnostics, ACD Bio, Newark, CA), and the RNAscope**®** LS 2.5 Probe-Mm-Ackr1 (ACD Bio, Catalog No: 450238). To confirm adequate RNA integrity of the FFPE samples, a medium-expressing housekeeping gene (peptidyl isomerase B, PPIB [ACD Bio]) was applied to a subset of SARS-CoV-2-infected and mock-infected samples as a positive control. A probe targeting the dapB gene (ACD Bio) was used as a negative control. The 5 µm samples were baked, deparaffinized, and underwent epitope retrieval using Leica Epitope Retrieval solution 2, pH 9.0 (Leica Microsystems) at 95°C for 15 min followed by a Protease III (ACD Bio) treatment at 40°C. Tissues then were hybridized with the *Ackr1* probe for 120 min and visualized chromogenically with the RNAscope**®** 2.5 LS Reagent Kit-RED.

### Statistical analysis

Statistical differences were analyzed using GraphPad Prism (GraphPad Software, La Jolla, CA) using the statistical tests indicated in each figure legend.

## Results

### Ackr1 is highly upregulated in the lungs of SARS-CoV-2 infected mice

We first analyzed the expression of cytokines, chemokines and chemokine receptors in lungs of C57BL/6J WT mice after intranasal infection with SARS-CoV-2. Clinical strains of SARS-CoV-2 were unable to use mouse ACE2 for entry or cause morbidity or mortality in WT mice. Therefore, for our experiments we used the mouse-adapted MA10 strain of the virus (Leist et al., 2020). JAX664 C57BL/6 mice of 4-6 months of age were infected intranasally with approximately 8.62 × 10^4^ TCID_50_. Both male and female mice demonstrated a similar extent of weight loss and mortality across independent experiments. Lung samples from infected and mock (PBS)-infected mice were harvested at 4 dpi and the expression of chemokine ligand and receptor genes was comprehensively analyzed by qPCR. Among the chemokine ligands, we found a significant upregulation of the monocyte-targeted chemokines *Ccl2*, *Ccl7* and *Ccl12*; the neutrophil-targeted chemokines *Cxcl1*, *Cxcl2* and *Cxcl3*; Type 1 immune response chemokines *Ccl3*, *Ccl4*, *Cxcl9*, *Cxcl10* and *Cxcl11*; *Cxcl12*, which exerts pleiotropic effects on lymphocytes; the B cell chemoattractant, *Cxcl13*; and *Xcl1*, which chemoattracts cross-presenting CD8^+^ dendritic cells (Fig. 1A) (Griffith et al., 2014; Majumdar and Murphy, 2021). Consistent with this, the cognate G protein-coupled receptors for many of these chemokines, *Ccr1*, *Ccr2*, *Ccr5* and *Cxcr2,* were also upregulated after infection. However, by far the most highly upregulated chemokine receptor after SARS-CoV-2 infection in these mice was *Ackr1 (Fig. 1B)*.

**Figure 1:**
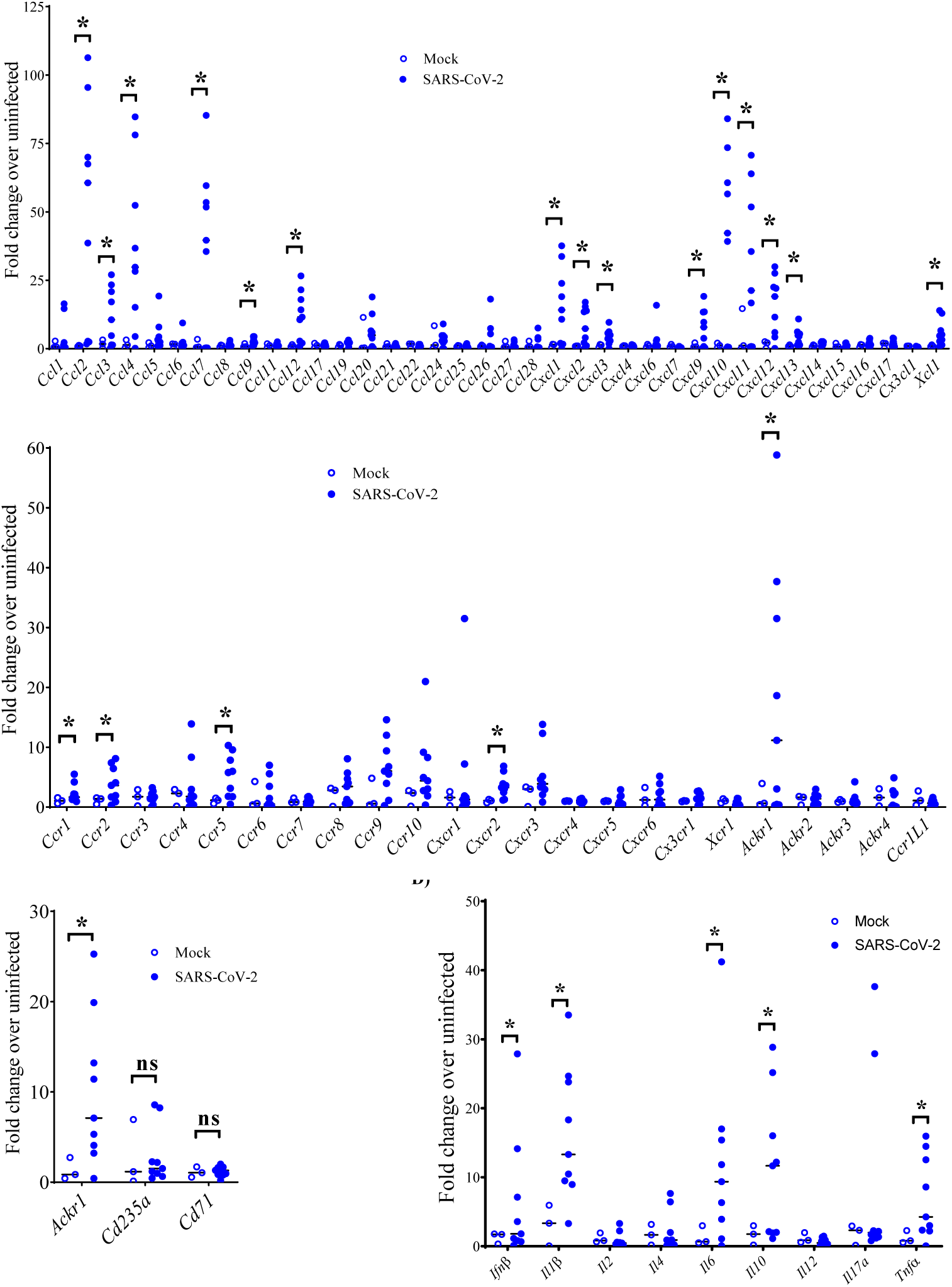
SARS-CoV-2 infection induces strong *Ackr1* expression in lung of WT mice. WT mice were intranasally infected with 8.62 × 10^4^ TCID_50_ of SARS-CoV-2 MA10, while mock-infected mice received PBS. After 4 days of infection, the mice were sacrificed, and lung tissues were harvested for gene expression studies by RT-PCR. **(A)** Chemokines, **(B)** chemokine receptors, **(C)** RBC maturation markers and **(D)** cytokines. The genes analyzed by RT-PCR are indicated on the x-axes. Data are from one experiment with 3-9 mice per group. Each dot corresponds to one mouse and the median value for each group is indicated with a black horizontal line. ns, not significant and *, p< 0.05, as determined by two-way ANOVA with Sidak’s multiple comparison test for all the data sets.

Since severe COVID-19 is a major risk factor for microvascular clots and pulmonary embolism (Katsoularis et al., 2022; Poissy et al., 2020; Wichmann et al., 2020), and ACKR1 is known to be expressed on red blood cells (RBCs), and since red cells do contain mRNA (Kabanova et al., 2009), we next studied whether increased *Ackr1* gene expression might be due to *Ackr1* transcripts accumulated in lung blood clots of infected animals. We found that mRNA for the mature erythrocyte markers, *Cd235a* and *Cd71*, was not increased at 4 dpi with SARS-CoV-2 infection (Fig. 1C), indicating that RBC were unlikely to be the source of increased *Ackr1* transcripts in the infected lung. We also observed upregulation of *Il10* mRNA as well as mRNA for the pro-inflammatory cytokines *Ifnβ*, *Il1β*, *Il6* and *Tnfα,* consistent with previously published data (Winkler et al., 2020) (Fig. 1D).

To investigate the kinetics of *Ackr1* gene expression, we harvested lung tissues from 1, 4 and 6 dpi with SARS-CoV-2 along with tissue from mock-infected mice. *Ackr1* mRNA was upregulated as early as day 1 after infection, and peaked by 4 dpi (Fig. 2A, Fig. 3A). On day 6 *Ackr1* mRNA levels slightly waned; nonetheless, they were still higher than for all other receptors studied (Fig. 2A). The expression of the pro-inflammatory genes, *Il6* and *Ifnβ* remained high till 6 dpi (Fig. 2B). Although not statistically significant (p = 0.201) and to a lesser degree than in C57BL/6 mice, SARS-CoV-2 infection appeared to also induce *Ackr1* expression in BALB/c mice at 4 dpi (Fig. 3B). Intranasal infection with MCMV and Influenza A Virus (IAV) strain PR8 also induced *Ackr1* mRNA in the lung of C57BL/6 mice; however, the fold-induction levels were considerably lower compared to those observed after SARS-CoV-2 infection (Fig. 3C).

**Figure 2:**
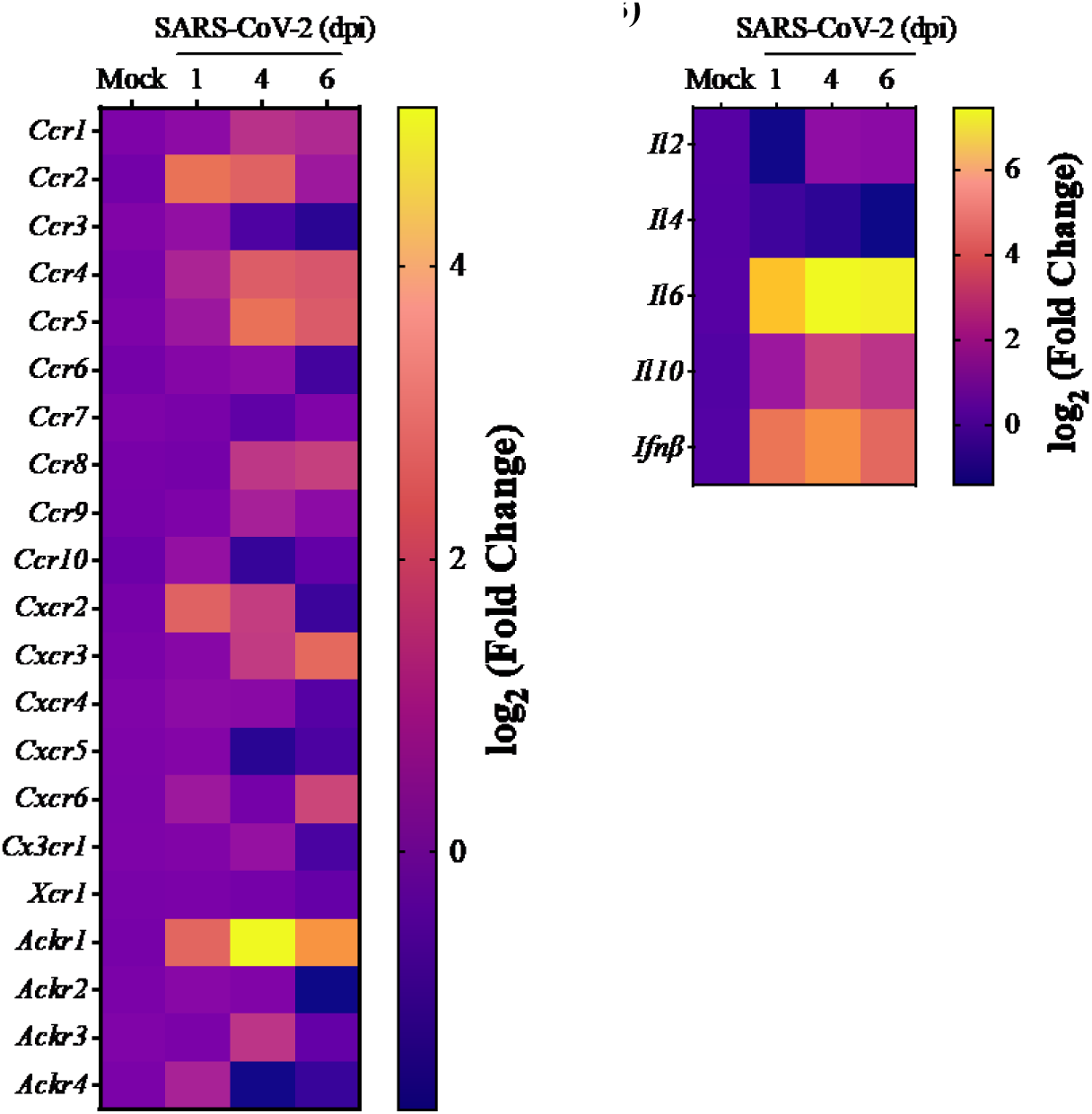
*Ackr1* mRNA is rapidly induced and shows sustained expression in the lung of SARS-CoV-2-infected mice. WT C57BL/6 mice were intranasally infected with 8.62 × 10^4^ TCID_50_ of SARS-CoV-2 MA10. After 1, 4 and 6 dpi (days post infection), the lung tissues were harvested along with lung tissues from mock-infected mice. **(A and B)** Heat map visualizing the expression of chemokine receptor genes and cytokine genes after SARS-CoV-2 infection, as measured by RT-PCR. The fold change for individual genes was calculated for SARS-CoV-2 infected mice relative to the level of expression in mock-infected mice sacrificed on the same day. Data were derived from one experiment consisting of 2-6 mice per experimental group.

**Figure 3:**
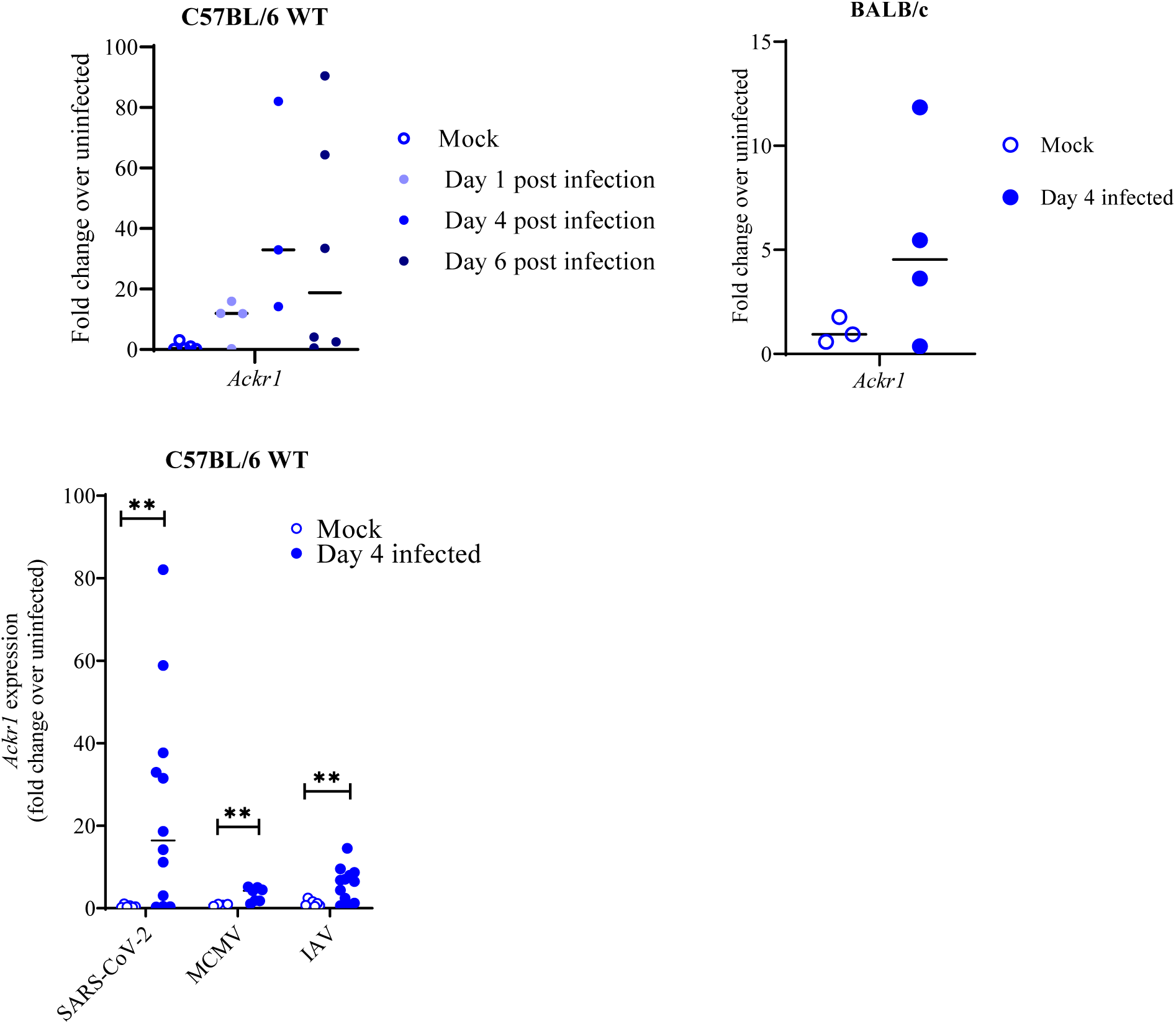
*Ackr1* mRNA induction in mouse lung is observed early after SARS-CoV-2 infection and is greater than after MCMV and IAV infection. The mice were intranasally infected with 8.62 × 10^4^ TCID_50_ of SARS-CoV-2 MA10, while the mock-infected mice were administered PBS. At the indicted timepoints, the mice were sacrificed, and lung expression of *Ackr1* was quantified by RT-PCR. **(A)** C57BL/6 JAX664 WT mice. **(B)** BALB/c mice. (C) *Ackr1* expression in lung of SARS-CoV-2, MCMV and IAV-infected mice. C57BL/6 JAX664 WT mice were intranasally infected with 8.62 × 10^4^ TCID_50_ of SARS-CoV-2 MA10, 10^5^ pfu of MCMV and 100 TCID_50_ of IAV. Data are from one experiment with 3-12 mice per group. Each dot corresponds to data from one mouse. The median value is indicated by a black horizontal line. **, p< 0.01, as determined by multiple unpaired t test.

### Ackr1 expression is restricted to venule and arteriole endothelium in SARS-CoV-2 infected mouse lung

At 4 dpi with SARS-CoV-2, at the peak of *Ackr1* pulmonary expression, lung was harvested from challenged mice and analyzed to determine *Ackr1* mRNA localization. At the site of a representative pulmonary arteriole with mild mononuclear perivascular inflammation (Fig. 4A), SARS-CoV-2 spike protein antigen was found within alveolar epithelial cells and alveolar or perivascular mononuclear cells (a finding consistent with previous published MA SARS-CoV-2 studies) (Leist et al., 2020), but not in smooth muscle or endothelial cells (ECs) of the arteriole (Fig. 4B). CD31 staining clearly highlighted the endothelium of the arteriole and the alveolar capillaries (Fig. 4C). Notably, *Ackr1* mRNA expression, as evidenced by the intense red positive hybridization signal, was associated with the endothelium of some but not all small to medium-sized vessels in SARS-CoV-2-infected mice at day 4 post-infection (Fig. 4D, Fig. 5A). Furthermore, in affected arterioles, *Ackr1* mRNA was only found in a subset of the arterial ECs (Fig. 4D) and not anywhere outside the arteriole and did not co-localize with SARS-CoV-2-positive cells. Expression of *Ackr1* mRNA was restricted to vascular endothelium, both venular and arterial (Fig. 5A and B). Additionally, positive hybridization signal was not observed in bronchiolar epithelial cells or in intra-airway inflammatory or desquamated bronchiolar epithelial cells of infected mice. Chromagenic *in situ* hybridization for *Ackr1* mRNA was specific and distinct from background (Fig. 5C and D) and was completely absent on vascular endothelium in the lungs of mock-infected mice (Fig. 5E). Therefore, in agreement with the qPCR data (Fig. 1B), a strong induction of *Ackr1* expression after infection was observed.

**Figure 4:**
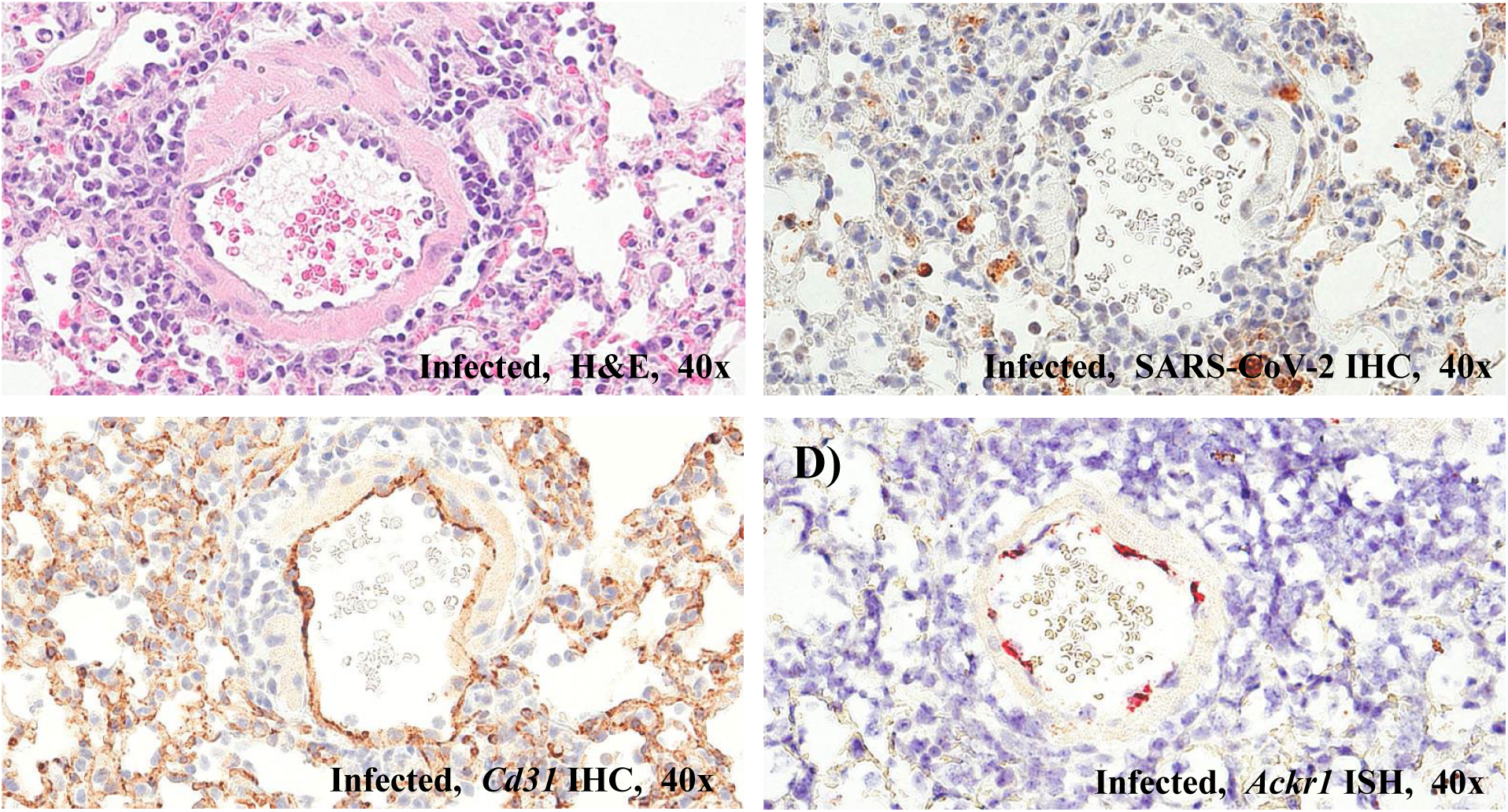
*Ackr1* mRNA is expressed by arteriolar endothelium in SARS-CoV-2 infected mouse lung at 4 dpi. **(A-D)** Serial sections of FFPE lung tissue from 4-month-old C57BL/6J WT mice that were intranasally infected with 8.62 × 10^4^ TCID_50_ of SARS-CoV-2 MA10 and harvested 4 dpi. **(A)** H&E. A pulmonary arteriole with mild mononuclear perivascular inflammation and reactive ECs. 40x. **(B)** SARS-CoV-2 IHC. SARS-CoV-2 viral antigen is not associated with the endothelium of the pulmonary vessel. Adjacent to the vessel, viral antigen (brown) is present in low numbers of alveolar mononuclear cells and alveolar epithelial cells. 40x. **(C)** CD31 IHC (ECs). Serial section of the same vessel. Strong positive CD31 immunostaining (brown) of ECs lining an arteriole as well as alveolar septae (alveolar capillary ECs). 40x. **(D)** RNAscope^®^ *Ackr1* chromagenic ISH staining. Positive hybridization signal (red) is observed in cells along the endothelium of the pulmonary arteriole. 40x.

**Figure. 5:**
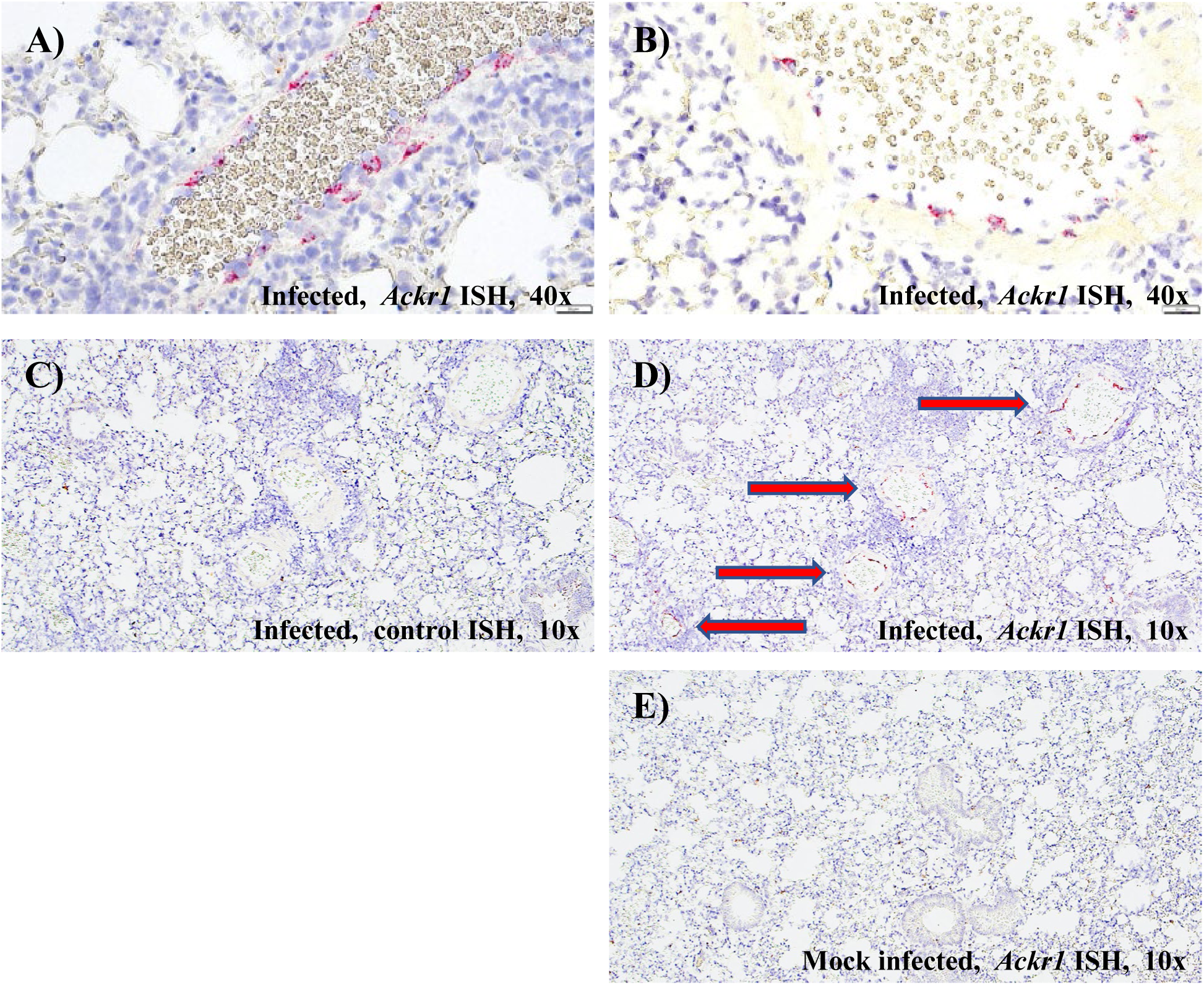
*Ackr1* mRNA expression is induced in pulmonary venules after SARS-CoV-2 infection. Serial sections of FFPE lung tissue from 4-month-old C57BL/6J WT mice that were intranasally infected with 8.62 × 10^4^ TCID_50_ of SARS-CoV-2 MA10 and harvested 4 dpi. **(A-B)** RNAscope^®^ *Ackr1* chromagenic ISH staining, 40x. Positive hybridization signal (red) was observed in the endothelial cells (EC) lining some but not all small to medium sized vessels. The intracellular signal varied from low (few dots) to high characterized by filling of the entire spindled cytoplasmic space. Not all ECs in affected vessels expressed signal. Positive hybridization signal was not readily observed in alveolar septa (no signal associated with alveolar capillary ECs). Signal was not observed in bronchiolar epithelial cells nor in intra-airway inflammation or desquamated bronchiolar epithelial cells (bronchioles are not present in the images). **(C)** RNAscope^®^ ISH with a negative-control target probe targeting the bacterial *dapB* gene, 10x. **(D)** RNAscope^®^ *Ackr1* chromagenic ISH staining. Positive hybridization signal (red arrows) was observed along the endothelium of multiple pulmonary vessels, 10x. **(E)** Section of FFPE lung tissue from mock-infected (PBS vehicle) 4-month-old C57BL/6J WT mice. RNAscope^®^ *Ackr1* chromagenic ISH staining, 10x. No hybridization signal was observed in this section of lung from a mock-infected mouse.

### Ackr1^-/-^ mice are resistant to SARS-CoV-2-induced mortality

To determine whether ACKR1 is functionally relevant in SARS-CoV-2 infection, we infected *Ackr1*^-/-^ mice along with other mouse strains deficient for genes regulating chemotaxis, as well as non-littermate WT C57BL/6J mice, which were used as controls, with a dose equivalent to 8.62 × 10^4^ TCID_50_ of SARS-CoV-2 MA10 and monitored weight loss and mortality over the course of 14 days. In WT control mice, this viral dose corresponded to approximately the LD_60_, which enabled detection of genes that might either improve or worsen outcome in the model. Mice that lost 25% of their initial body weight were euthanized according to animal health and safety welfare protocol guidelines. Only the *Ackr1*^-/-^ mice had lower mortality than the reference WT mice (*p*=0.0014, Mantel-Cox test). Mortality in infected *Cxcr3*^-/-^ and *Fpr1/2*^-/-^ mice was equivalent to that in WT mice, whereas mortality in infected *Ackr2*^-/-^, *Ccr5*^-/-^, *Ccr6*^-/-^ and *Camp*^-/-^ mice was slightly higher than in WT control mice (Fig. 6A). The protection conferred by ACKR1 deficiency was stronger in male mice than female mice and did not reach statistical significance in female mice (Fig. 6B and C). *Ackr1* upregulation in lungs of WT male mice trended higher than in lungs of female WT mice (Fig. S1). Weight loss in *Ackr1*^-/-^ mice was similar to that of WT mice, among both survivors and non-survivors (Fig. 6D-F).

**Figure 6:**
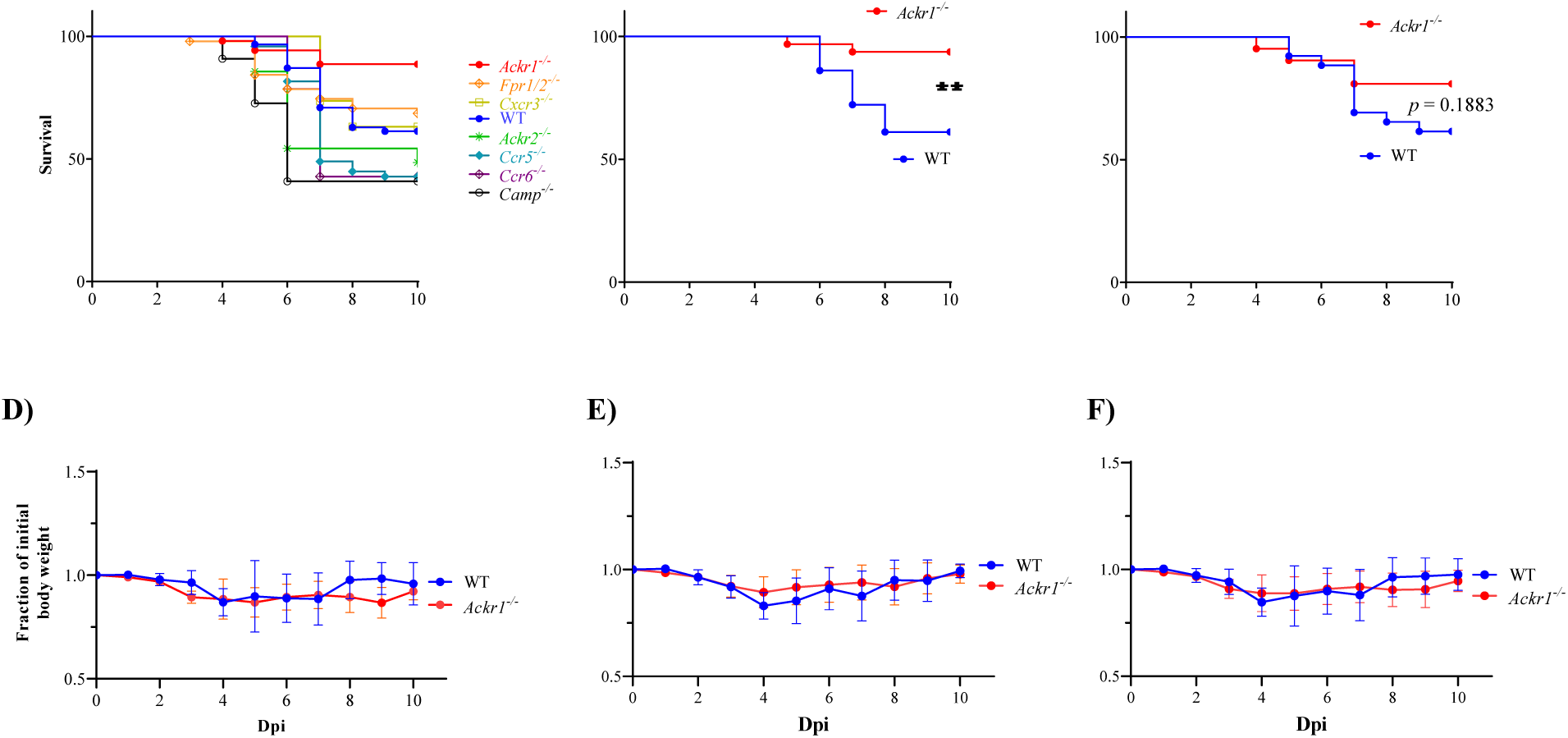
ACKR1 deficiency in mice confers protection towards SARS-CoV-2-induced mortality. 4-6-month-old mice were intranasally infected with 8.62 × 10^4^ TCID_50_ of MA SARS-CoV-2. **(A-C)** Survival. Mortality was recorded over the course of 14 days post-infection (dpi). Mice that lost 25% of their initial body weights (endpoint) were euthanized according to predefined guidelines. Mouse genotypes are color-coded as indicated on the right side of the figure. **(A)** Survival of combined male and female mice infected with MA SARS-CoV-2. **(B-C)** Data from panel A stratified by sex. **(D-F)** Body weight after infection. **(D)** Combined male and female mice. **(E-F)** Data from panel **D** stratified by sex. **E**, male; **F**, female. Cumulative data are summarized from a minimum of 5 independent experiments, n=14-62 mice/genotype as indicated in panels A-C. Statistical significance in **(A-C)** were assessed by Mantel-Cox test. **, *p*< 0.01.

## Discussion

In the present study, we found that total genetic *Ackr1* deficiency increased survival in C57BL/6 mice infected with a lethal dose of a mouse-adapted strain of SARS-CoV-2. Males benefited more than females. Unlike G protein-coupled chemokine receptors, atypical chemokine receptors do not activate G proteins and are expressed in atypical locations. ACKR1 is the most promiscuous of all chemokine receptors, binding with high affinity to many but not all inflammatory chemokines from both the CC and CXC subclasses and is not expressed on leukocytes. Instead in addition to erythrocytes, it has been detected on post-capillary venular ECs (Lee et al., 2003; Peiper et al., 1995; Hadley et al., 1994), including in the lung (Schupp et al., 2021), as well as systemic venous ECs (De Rooij et al., 2023) and Purkinje cells of the cerebellum (Horuk et al., 1996). To our knowledge, our study is the first to define *Ackr1* expression in arteriolar endothelial cells and induction by SARS-CoV-2 (Fig. 4 and 5).

*ACKR1* is considered a marker gene for systemic venous ECs (De Rooij et al., 2023). Previous studies have demonstrated that expansion of ACKR1^+^ venous ECs occurs in aged mice when persistent fibrosis is induced by bleomycin (Raslan et al., 2023). These ACKR1^+^ venous ECs are closely associated with alveolar regions showing reduced capillary density and alveolar thickening. In addition, fibrotic tissue from patients with idiopathic pulmonary fibrosis have been reported to be populated by ACKR1^+^ venous ECs (Raslan et al., 2023). Single-nucleus RNA-sequencing studies revealed increased abundance of systemic venous ECs in lungs of COVID-19 decedents and idiopathic pulmonary fibrosis patients compared to lung tissues from healthy controls. Our results validate restriction of *ACKR1* expression to venular and arterial ECs in SARS-CoV-2-infected mouse lung (Fig. 5).

Our result aligns with exceptionally low COVID-19 mortality rates in sub-Saharan Africa (Cabore et al., 2022), a unique geographic region in which selective genetic ACKR1 deficiency in erythrocytes is fixed in the population (Fig. 7A, B) (Howes et al., 2011). While the frequencies of confirmed infection (cases/100,000) differed significantly across regions with higher prevalence of ACKR1 deficiency in erythrocytes [552.42 (Ghana), 129.37 (Nigeria), 334.84 (Côte d’Ivoire), 2200.67 (Gabon) and 97.31 (Sierra Leone)] compared to regions with low prevalence of ACKR1 deficiency in erythrocytes [9752.51 (Tunisia), 7382.16 (Libya), 504.2 (Egypt), 6866.67 (South Africa) and 6738.62 (Namibia)], a seroprevalence study estimated that the actual prevalence may vastly differ among African countries, with Sierra Leone having a seroprevalence level 43-times higher than reported infections, and South Africa having a seroprevalence estimate marginally higher than reported (47.4% vs. 35.6-41.2%) (Cabore et al., 2022). Further, in a 6-month prospective study in 2020 in the West African country of Mali, the cumulative SARS-CoV-2 seroconversion rate was 58.5%, yet none of 2657 individuals died of COVID-19 and only 3 were hospitalized (Sagara et al., 2022). In contrast, African Americans have a higher COVID-19 mortality rate than sub-Saharan Africans; however, the red cell *ACKR1*^-/-^ genotype frequency is lower and comorbidity rates may be higher in African Americans compared to sub-Saharan Africans.

**Figure 7:**
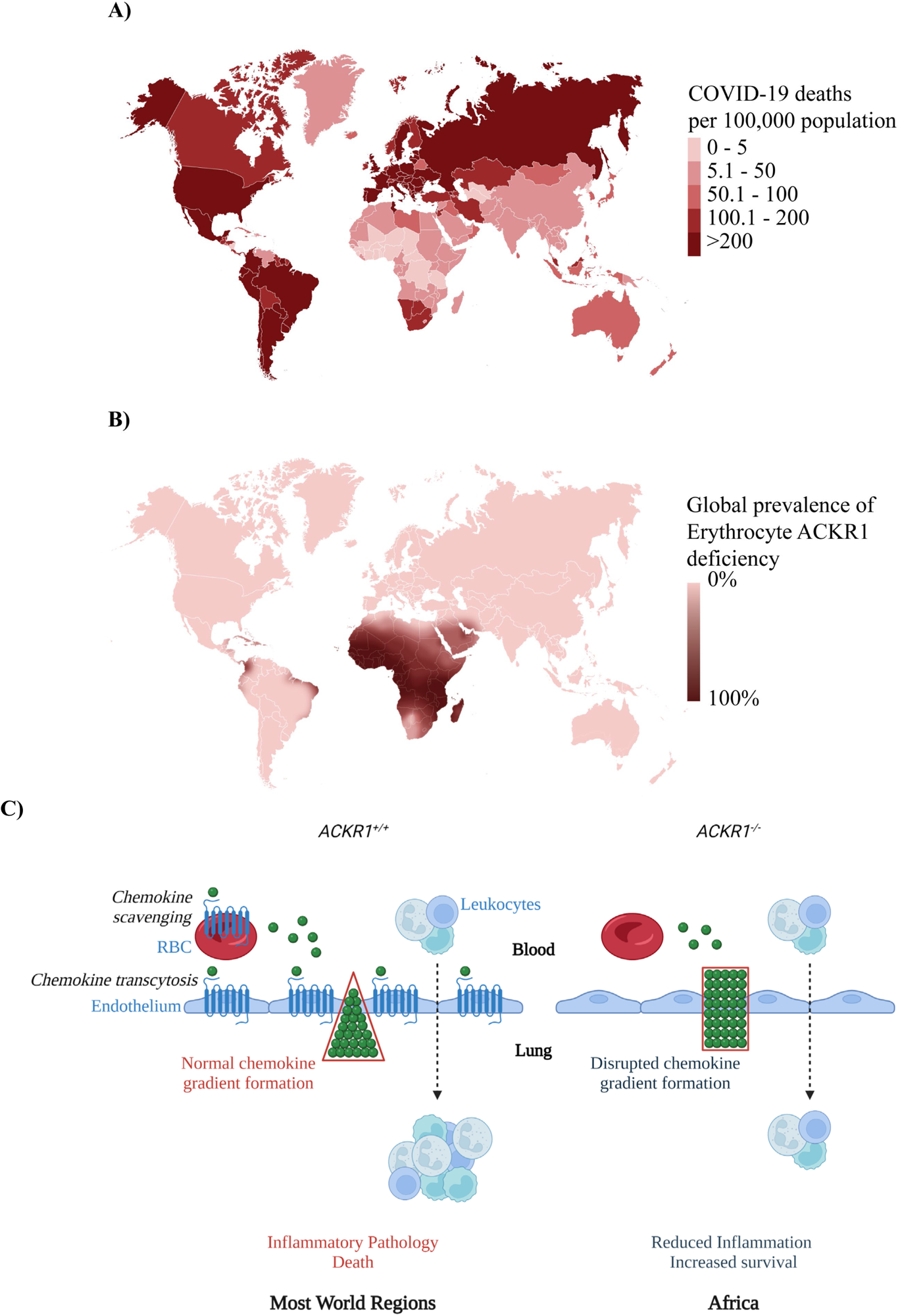
ACKR1 deficiency and SARS-COV-2 pathogenesis. **(A)** COVID-19 deaths per 100,000 population (adapted from https://covid19.who.int/). **(B)** Global distribution of erythrocyte ACKR1 deficiency (adapted from Howes *et al*., 2011). **(C)** Putative role of ACKR1 during SARS-CoV-2 pathogenesis. ACKR1 is expressed on RBCs as well as ECs to regulate the gradient of chemokines by scavenging and transcytosis, respectively. During severe cases of SARS-CoV-2, cytokine storm causes upregulation of inflammatory chemokines in circulation and lung tissue. Chemokine gradient formation may mediate infiltration of lung by leukocytes that initiate development and progression of ARDS-linked inflammation and death. Absence of ACKR1 affects scavenging and transcytosis of chemokines that in turn may influence the shape of chemokine gradients and consequently the infiltration of leukocytes within the SARS-CoV-2-infected lung, thereby negatively regulating ARDS development, and improving survival. The majority of the population in sub-Saharan Africa lacks ACKR1 expression on RBCs, which might confer protection from severe COVID-19.

In humans, erythrocyte-specific *ACKR1* deficiency is caused by a single nucleotide change in the *ACKR1* promoter (rs2814778) (Höher et al., 2017), which interferes with binding of the erythroid-specific transcription factor GATA-1. Thus, this mutation does not affect expression on the other known ACKR1^+^ cell types, ECs and Purkinje cells of the cerebellum (Tournamille et al., 1995). Erythrocyte-specific *ACKR1* deficiency is thought to have been fixed in the sub-Saharan African population under the selective pressure of infection with the malarial parasite *Plasmodium vivax*, which uses ACKR1 as an entry factor to establish erythrocyte infection (Miller et al., 1976). Since we analyzed a total *Ackr1^-/-^*mouse, additional work will be needed, and is now underway, to define whether protection in the mouse model is conferred by erythrocyte or endothelial cell *Ackr1* deficiency, or both, and to determine whether human erythrocyte-specific *ACKR1* deficiency is causally related to low COVID-19 mortality rates in sub-Saharan Africa. Erythrocytic and EC ACKR1 may affect different parameters of anti-viral response such as chemokine sequestration and transcytosis, respectively, ultimately affecting survival (Fig. 7C). Interestingly, both surviving as well as non-surviving *Ackr1*^-/-^ mice lose similar proportions of body weight upon SARS-CoV-2 infection compared to WT mice (Fig. 6D-E). By day 7, viral replication is absent in lungs of MA10 SARS-CoV-2-infected WT mice, coinciding with recovery of body weight in surviving mice (Leist et al., 2020). We are currently assessing the effect of ACKR1 deficiency on viral load in infected mice.

On ECs ACKR1 can present tissue-derived chemokines on the luminal surface of vessels to rolling leukocytes via transcytosis, while in erythrocytes it can function as a blood reservoir for chemokine action or else as a chemokine sink (Bonecchi and Graham, 2016). Both erythrocyte and endothelial cell expression may function to shape chemokine gradients and promote leukocyte trafficking into infected or inflamed sites from the blood. Disruption of either function might disrupt gradients and inhibit inflammation (Lee et al., 2006). In this regard, plasma levels of the monocyte-attracting chemokine CCL2 and the neutrophil-attracting chemokine CXCL1 were reported to be ∼2-fold higher in ACKR1-negative volunteers. Conversely, erythrocyte-bound CCL2, CXCL8 and CXCL1 were significantly higher in LPS-infused ACKR1-positive individuals (Mayr et al., 2008). Moreover, almost all individuals with the red cell ACKR1-null phenotype have benign ethnic neutropenia, a term used to describe asymptomatic resting ANC levels <1500 cells/µL (Reich et al., 2009; Legge et al., 2020), which might also attenuate immunopathology in COVID-19. The ANC in *Ackr1*^-/-^ mice is normal but blood neutrophils from both *Ackr1*^-/-^ mice and ACKR1-negative humans express higher amounts of cell-surface CD16/32 and CD45 compared to WT neutrophils. CD16/32 is important for neutrophil activation by immune complexes, and CD45 amplifies CD16/32 function (Duchene et al., 2017). Given these precedents, since severe COVID-19 has been associated with ARDS in the context of viral clearance, ACKR1 deficiency may dampen immunopathology, possibly explaining the reduced mortality we observed in *Ackr1*^-/-^ mice (Fig. 6A-C, 7C). Additional work will be needed to address the protective mechanisms in infected *Ackr1*^-/-^ mice, including assessments of viral burden and clearance rates, the extent and type of immunopathology as well as chemokine levels in blood and tissue.

Unlike *Ackr1*^-/-^ mice, *Ackr2*^-/-^ mice were not resistant to SARS-CoV-2 lethality in our study, although they have been reported to be protected against IAV infection (Gowhari Shabgah et al., 2022). We also did not observe increased survival in *Ccr5*^-/-^ or *Ccr6*^-/-^ mice during SARS-CoV-2 infection. CCR5 blocking agents have been tested in severe COVID-19 patients with mixed results. In contrast to anecdotal reports of benefit with Leronlimab, a monoclonal antibody directed against CCR5, (Patterson et al., 2021; Yang et al., 2021), Cenicriviroc, a dual small molecule CCR2/5 antagonist, was ineffective (Files et al., 2023).

In conclusion, our study establishes a role for ACKR1 during SARS-CoV-2 pathogenesis in mice. The results provide a potential genetic mechanism to explain the apparent reduced mortality in ACKR1-negative individuals and potentially a new therapeutic target for severe COVID-19.

## Acknowledgments

We thank Dr. Ralph Baric for providing the MA10 of SARS-CoV-2. We would also like to thank Mr. Alexander Stewart and Mr. Ryan Kissinger for generating the maps in Fig. 7A and B. Fig. 7C was created in BioRender.com.

## Author Contributions

PMM conceptualized the study. SM, JDW, SMP, MM, SGC and TB performed the experiments. NL, RJ, XL and HZ provided the resources. SM, JDW, SMP, J-LG, BLK, JMF and DAA analyzed the data. All authors have read and agreed to the final version of the manuscript.

## List of abbreviations

ACKR1: Atypical chemokine receptor 1
ARDS: Acute respiratory distress syndrome
*CCR*: CC motif chemokine receptor
COVID-19: Coronavirus disease-2019
*CXCR*: C-X motif chemokine receptor
DPI: Days post-infection
ECs: Endothelial cells
FPR: Formyl peptide receptor
GWAS: Genome-wide association study
IAV: Influenza A virus
ISH: *In situ* Hybridization
KO: Knockout
MA: Mouse-adapted
MCMV: Murine cytomegalovirus
MERS-CoV: Middle East Respiratory Syndrome Coronavirus
RBCs: Red blood cells
SARS-CoV: Severe acute respiratory syndrome coronavirus
WT: Wild type

