## Supplementary Figure 1 for "*Atypical Chemokine Receptor 1 (Ackr1)*-deficient Mice Resist Lethal SARS-CoV-2 Challenge"

### Supplementary Material

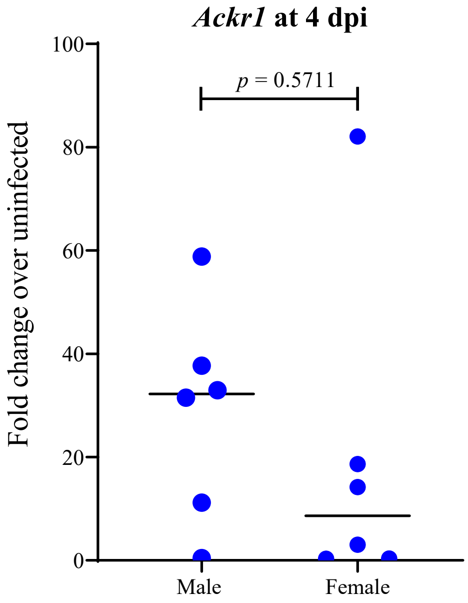


**Figure S1: *Ackr1* mRNA expression in lungs of SARS-CoV-2-infected mice stratified by sex.** WT mice were intranasally infected with SARS-CoV-2 or PBS as control. Four dpi, the mice were sacrificed, and lung tissues were harvested for *Ackr1* expression by qPCR. Sexes of the mice are indicated on the x-axes. Each dot corresponds to data from one mouse and the median value for each group is indicated with a horizontal black line. Data were analyzed by the two-tailed unpaired t test.
